# A fungal hemophore relay mediates heme transfer via transient protein interactions

**DOI:** 10.64898/2026.05.08.723727

**Authors:** Udita Roy, Elian Far, Soham Chattopadhyay, Ziva Weissman, Daniel Kornitzer

## Abstract

The fungal pathobiont *Candida albicans* acquires heme from host proteins via a set of soluble and anchored extracellular CFEM-type hemophores that can capture the bound heme and exchange it, eventually delivering it to the cell membrane for endocytosis into the yeast cell. Yet the molecular mechanism by which the heme is transferred through this protein cascade across the cell envelope remains unclear. To address this mechanism, we developed a set a fusions of three *C. albicans* hemophores with fluorescent proteins. Fluorescence of these fusion proteins is strongly quenched when heme is bound to the hemophore moiety, enabling to measure heme transfer instantaneously. Kinetic analysis of the different transfer reactions reveals that heme transfer from the host protein to the CFEM hemophores and between the CFEM hemophores are governed by different regimes. Kinetics of heme transfer from hemoglobin or serum albumin to the CFEM hemophores is mainly first-order, suggesting that heme stochastically released from host proteins is captured by the hemophores. In contrast, transfer of heme between the hemophores is near second-order, consistent with a mechanism requiring protein-protein interactions. To confirm this, we show that CFEM hemophores can interact in homodimeric and heterodimeric complexes. Furthermore, while dimerization-defective mutants of the soluble hemophore Csa2 are proficient in heme binding and extraction, they are defective in heme transfer. This supports a model of heme transfer by direct interaction between the members of the fungal hemophore cascade.

**SIGNIFICANCE:** Acquisition of extracellular heme as iron or heme sources is common in microorganisms, and particularly prevalent among pathogenic organisms that must contend with an iron-poor host environment. To extract heme from host proteins, microorganisms deploy various systems that include extracellular soluble and cell-anchored hemophores. Here we describe a new approach for monitoring heme binding and transfer in real time, based on the development of fluorescent derivatives of fungal hemophores. These novel reagents open a new window on the study of a common virulence factor of microbial pathogens.

## INTRODUCTION

Acquisition of essential trace metals can be challenging for microbes, due to scarcity or low solubility in the environment. Furthermore, micronutrients, and iron in particular, are also often sequestered by host organisms to limit pathogen proliferation (1, 2). One solution for this problem is the secretion by microbes of carrier molecules that extract essential ligands from the environment and deliver them to the cell. Examples of such molecules are siderophores (3), zincophores (4, 5) and hemophores (6), the latter of which enable microorganisms to utilize environmental heme as iron source in the absence of elemental iron. Known microbial hemophores include the HasA type in Gram(-) bacteria (7), the Isd-type hemophores in Gram(+) bacteria (8), the Rv0203 hemophore in *Mycobacterium tuberculosis* (9) and the CFEM domain hemophores in fungi (10, 11). In view of their importance and ubiquity, methods for directly monitoring ligand binding and release from such carriers would be useful. While siderophores are usually small molecules, the known classes of hemophores are all proteins, which makes them potentially more amenable to modification as reporters.

Fungal CFEM-type hemophores were identified in the pathobiont *C. albicans*, a normal member of the human microbiota that can cause severe invasive disease in immunocompromised patients (12–14). *C. albicans* can utilize heme from hemoglobin, possibly aided by its hemolytic activity (15, 16), and it can also utilize heme bound to serum albumin, a known scavenger of free heme in serum (17, 18). The *C. albicans* hemophore system includes a secreted protein (Csa2), a cell wall-anchored protein (Rbt5) and a plasma membrane-anchored protein (Pga7) (10, 11, 19). Crystal structure of Csa2 indicated that CFEM hemophores bind heme on a flat platform on the surface of the protein, and that heme-iron coordination depends on an unusual Asp residue that makes the binding ferric heme-specific (11). The CFEM hemophores are all able to extract heme from host heme-binding proteins such as hemoglobin and serum albumin, and are able to exchange it *in vitro*, suggesting that they form a cascade of proteins that transfer heme across the cell wall to the cell membrane, where it is delivered to the endocytic pathway (reviewed in (20)). At the plasma membrane, the ferric reductase-like proteins Frp1, which collaborates with Pga7, and its paralog Frp2, are involved in the internalization of the heme into the cell (21). Frp1 and Frp2 were found to contribute to the cellular heme reductase activity, and might therefore function in releasing the heme from the ferric heme-specific CFEM hemophores (11, 22).

While the outline of the heme transfer pathway in *Candida* is known, the details of the transfer reactions of heme from the host proteins to the CFEM hemophores, and from one type of hemophore to the next, are not well understood. Transfer mechanisms could involve capture of the ligand without interaction between proteins, or protein-protein interaction-mediated transfer. In bacterial systems, heme transfer between the Gram(-) *Serratia marcescens* HasA-HasR hemophores system and within the Gram(+) staphylococcal Isd hemophore cascade was shown to involve direct protein interactions (23–25). Regarding heme extraction from host proteins such as hemoglobin, the staphylococcal IsdB hemophore was shown to bind hemoglobin in order to extract its heme (26, 27), whereas heme capture from hemoglobin by the *Pseudomonas aeruginosa* HasA hemophore appears to involve passive capture of heme released from the globin (28).

To address the heme transfer mechanism in the *Candida spp*. hemophore cascade, we designed a set of CFEM hemophores tagged with fluorescent proteins (FPs) to serve as heme binding reporters. Proximity of the heme to the FP leads to quenching of the fluorescence, enabling measurement of the kinetics of heme transfer reactions. Analysis of the kinetics of the various reactions revealed different mechanisms for the host protein – hemophore and hemophore – hemophore reactions, and led to the identification of a protein-protein interaction surface on the CFEM hemophores.

## RESULTS

### Fluorescence quenching by heme of FP-CFEM hemophore fusion proteins

A cytochrome b_562_ heme-binding domain embedded in the Green Fluorescent Protein (GFP) has been shown to cause quenching of the fluorescence signal with increasing heme binding, presumably via fluorescence resonance energy transfer from the fluorophore to the heme (29–31). We attempted to take advantage of this quenching effect in order to monitor heme binding in real time to the *C. albicans* CFEM hemophore family proteins. To that end, we designed fusion proteins between Rbt5, Pga7 and Csa2 on the one hand, and the fluorescent proteins GFP-gamma (32) or mScarlet (33) on the other, and tested the effect of heme binding on FP fluorescence.

Rbt5 and Pga7 have a heme-binding CFEM domain at their N-terminus, followed by a serine/threonine-rich domain that is probably heavily O-mannosylated, and a predicted GPI anchor site (20). We tested insertions of the fluorescent proteins at two sites for Rbt5 and Pga7: either N-terminal to the CFEM domain (FP-CFEM), between the predicted signal peptide and the CFEM domain, or C-terminal to it, between the CFEM and Ser/Thr-rich domains (CFEM-FP). For Csa2, GFP was fused only N-terminally to the CFEM domain. The fusion proteins were expressed as recombinant proteins in *Pichia pastoris* (see Fig. 1 for a schematic description of the fusion proteins).

**Fig. 1.**
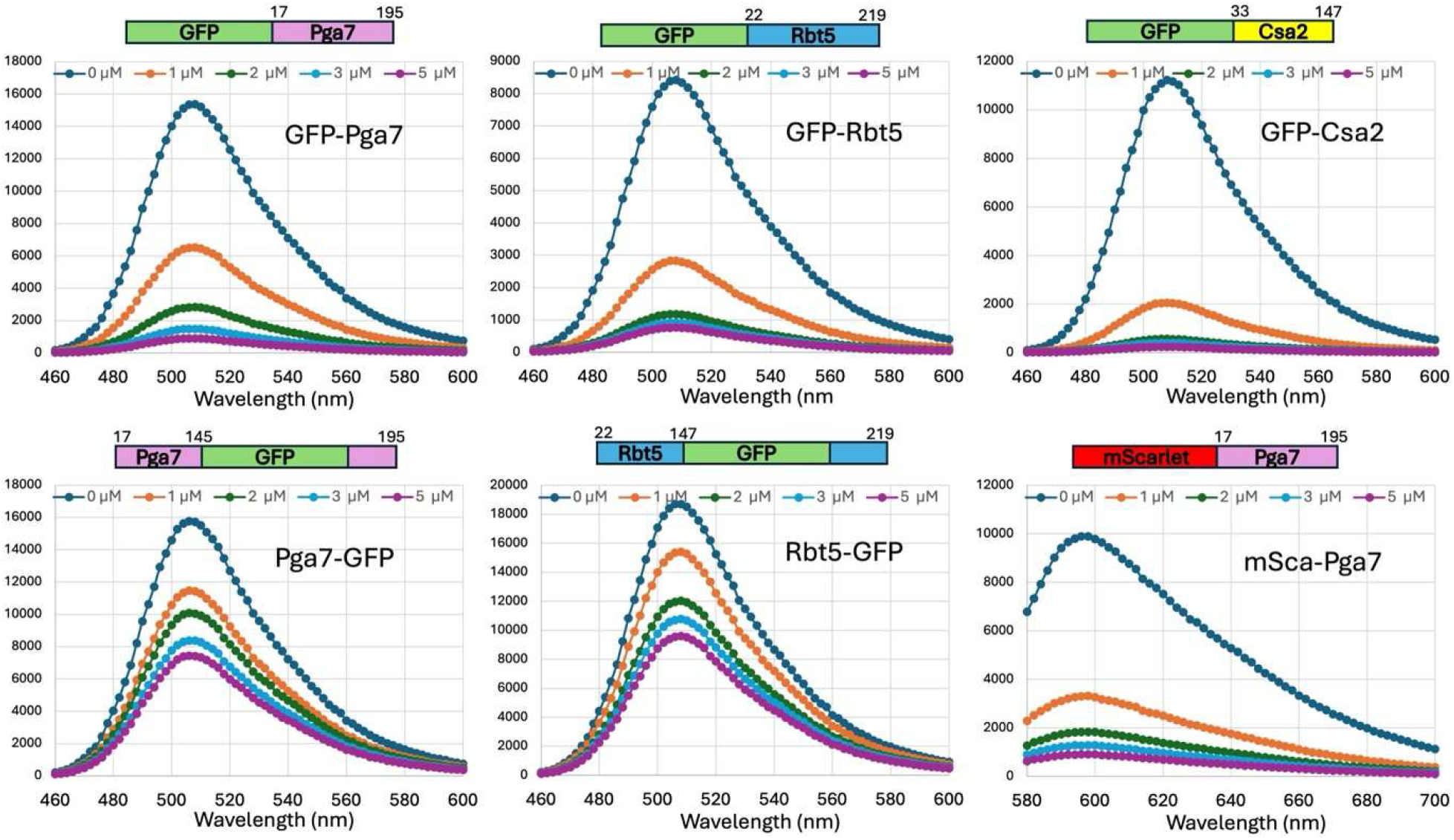
Heme binding quenches the fluorescence of fluorescent protein-CFEM hemophore fusions. Fluorescence emission scans of the different fusion proteins in the absence and presence of heme. 1 µM of each protein was mixed with the indicated concentrations of hemin, and scanned for fluorescence emission at an excitation wavelength of 420 nm (GFP) or 550 nm (mScarlet). The schematic structure of each fusion protein is shown above each graph with the hemophore amino acid coordinates.

Since the N-terminal fusions put the FP adjacent to the heme binding sites of the CFEM hemophores, we tested the ability of these proteins to bind heme by conventional visible light spectroscopy. All four fusion proteins bound heme like native CFEM hemophores, based on the shift of the main heme absorbance peak from 380-390 to a sharp Soret peak at 406 nm (11, 19) (Fig. S1).

We next tested the effect of heme addition on the fluorescence of the fusion proteins. To test the fluorescence, an emission scan was performed with fixed excitation in the absence or presence of increasing heme concentrations. Whereas the C-terminal fusions Pga7-GFP and Rbt5-GFP were quenched up to 2-fold, the N-terminal fusions GFP-Pga7, GFP-Rbt5 and mSca-Pga7 could be quenched over 10-fold, and GFP-Csa2 over 20-fold by the addition of heme (Fig. 1).

### Kinetics of heme transfer between hemoglobin or heme-albumin and the FP-tagged CFEM hemophores

CFEM hemophores are able to extract heme from hemoglobin or from human serum albumin (HSA) (11, 18, 19). We attempted to measure the transfer kinetics of heme from the host proteins to the fungal hemophores by mixing hemoglobin or heme-HSA with the FP-tagged hemophores GFP-Pga7, GFP-Rbt5, GFP-Csa2 and mScarlet-Pga7 and monitoring fluorescence.

Equimolar amounts of hemoglobin or heme-HSA and the FP-hemophore fusions were mixed at different concentrations, and fluorescence quenching was monitored at 0.2 sec intervals. As expected, exposure of the FP-hemophores to the host heme proteins resulted in a decrease in fluorescence, indicating transfer of heme to the hemophores. Initial reaction velocity was calculated and graphed against the starting concentration, and the data were fitted to a power equation, to obtain information about the kinetic reaction order, and gain insight into the transfer mechanism. As shown in Fig. 2A, the reaction kinetics with hemoglobin gave an exponent close to 1, indicating first-order behavior under these conditions. This suggests that the transfer mechanism does not involve a direct interaction between the hemophores and hemoglobin. The same reaction with holo-HSA as hemin donor, shown in Fig. 2B, resulted in a similar reaction rate, however fitting the rates to a power equation gave a slightly higher exponent of around 1.2 +/- 0.05. With both heme donors and all four FP-hemophore types, the heme transfer velocity was similar, between 0.05 – 0.1 nM / s at 1μM protein concentration.

**Fig. 2.**
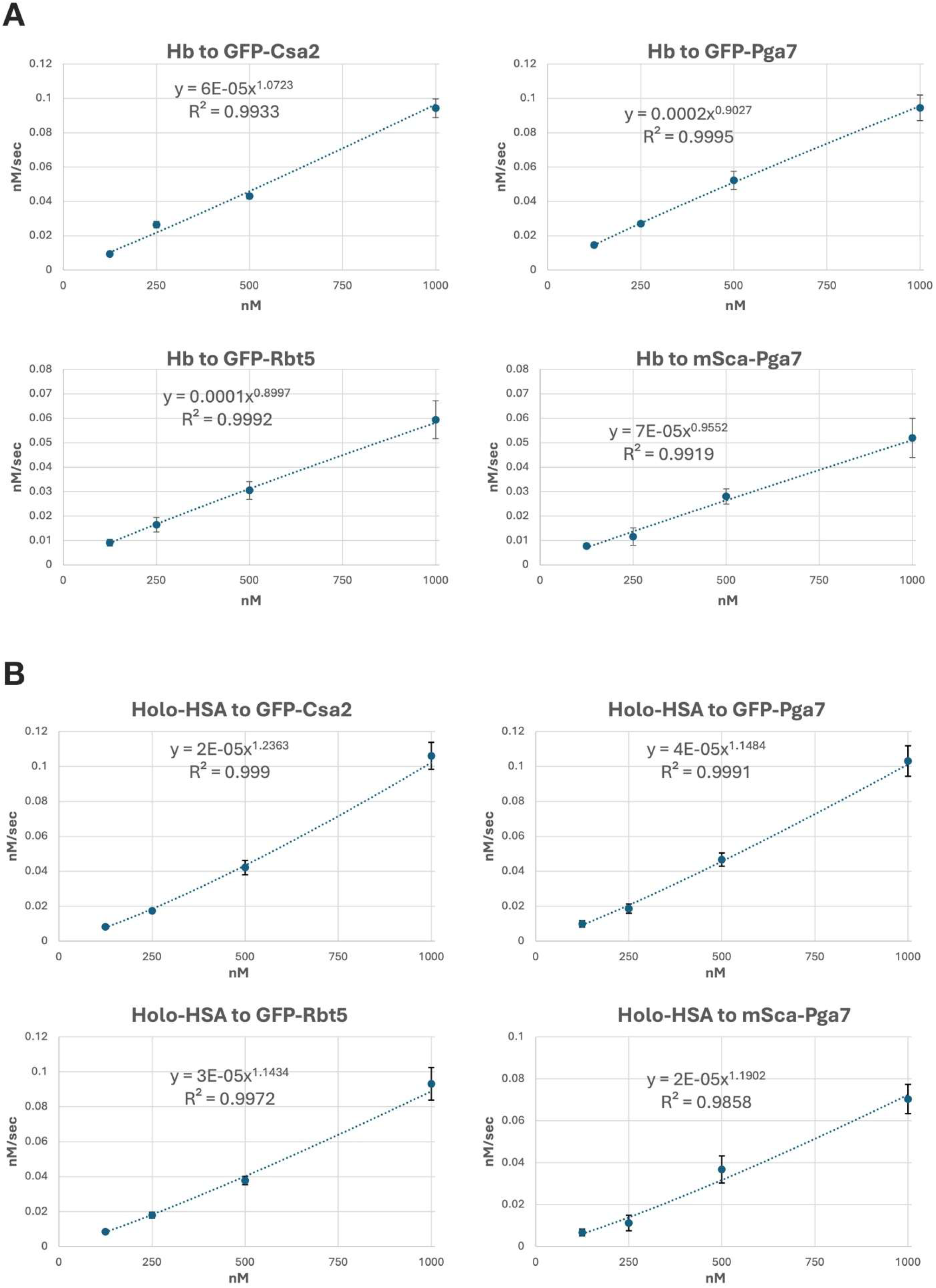
Heme transfer from hemoglobin and holo-HSA to the CFEM hemophores. The indicated apo-hemophores were mixed at equimolar concentration (125 nM, 250 nM, 500 nM, 1000 nM) with (A) bovine hemoglobin or (B) HSA loaded with hemin, and the fluorescence was monitored every 0.2 sec after injection. The fluorescence was converted to protein concentration, and for each datapoint, the average initial velocity for 5 to 8 injections is indicated (calculated from the rate in the first ∼ 5 sec, corresponding to about 10% of the total fluorescence decrease). The error bars indicate the standard error between the repeats.

### Kinetics of heme transfer between the CFEM hemophores

The current method to monitor heme transfer between CFEM proteins, namely mixing of apo- and holo-protein preparations followed by separation on a size exclusion column (11, 19) allowed to measure the equilibrium distribution but not the kinetics of transfer. Therefore, we took advantage of the quenching effect of heme to monitor its transfer between GFP-tagged and -untagged proteins, in order to measure transfer kinetics in real time.

We started with a fully heme-loaded GFP-CFEM hemophore fusion protein (holo-GFP-CFEM) and mixed it with the same concentration of a native apo-hemophore of the same or different kind. Reaction kinetics were followed at different starting concentrations at 0.05 sec intervals. The three native hemophores, Csa2, Rbt5 and Pga7 and their GFP fusions were tested in all combinations (Fig. 3). The three homotypic and six heterotypic mixtures showed much more rapid heme transfer than that from heme proteins to the hemophores (Fig. 2). When the data were fitted to a power equation, for most reactions, transfer velocity depended on the square or close to the square of the protein concentration, corresponding to second-order reactions. This is consistent with a substantial contribution of protein-protein interactions in the heme transfer reaction between hemophores.

**Fig. 3.**
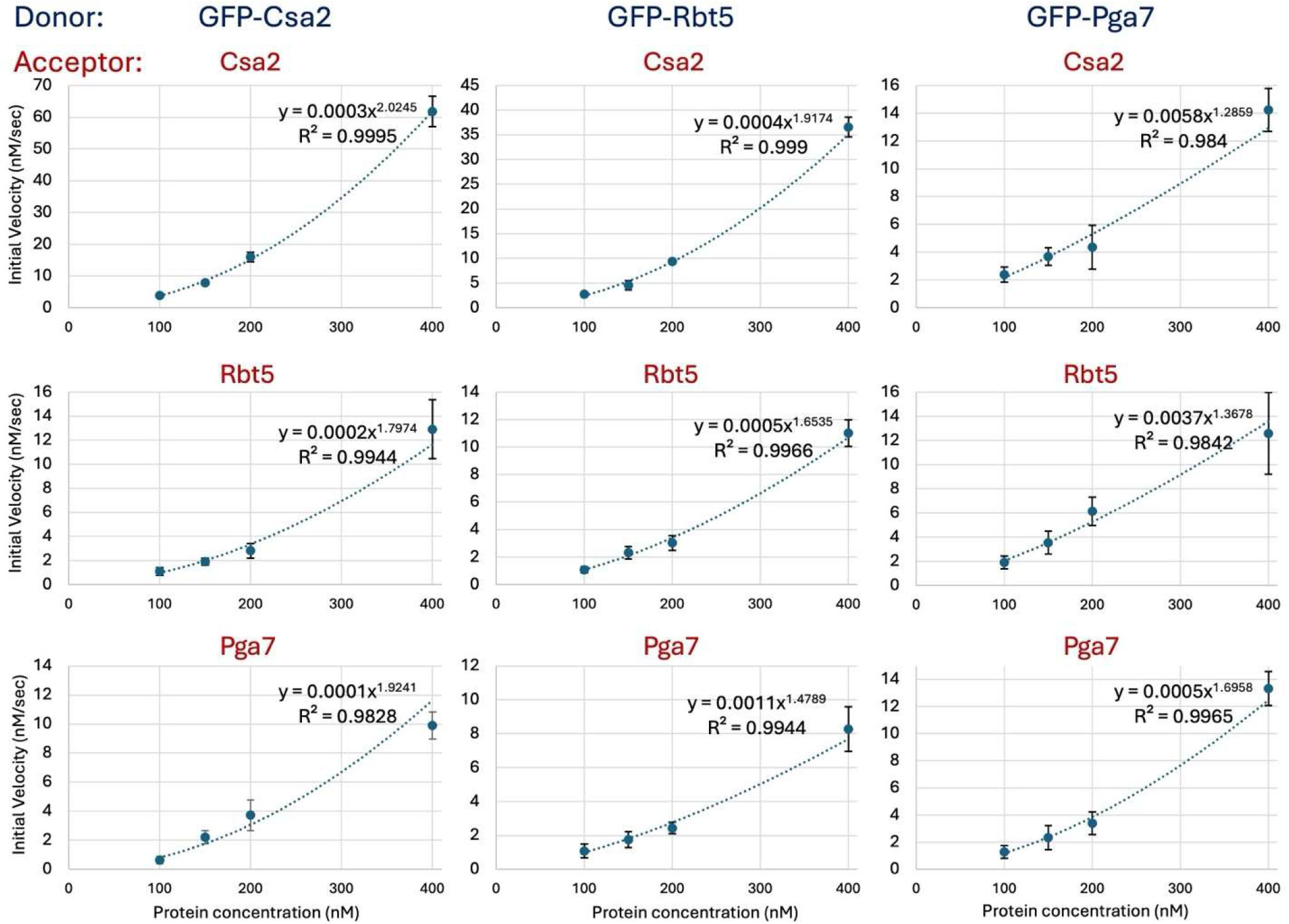
Kinetics of heme transfer between tagged holo-hemophores and untagged apo-hemophores. Combinations of each of the three GFP-tagged holo-hemophores with each of the three untagged hemophores were mixed, and de-quenching of the fluorescence by heme transfer was monitored with time at different starting protein concentrations. Each data point is the mean of 5-8 injections, with the error bars indicating the standard error between the repeats.

### CFEM hemophore dimerization

SEC-MALS analysis of apo- and holo-Csa2 had suggested that at high concentration, Csa2 is found at least partially in a dimer (11). To analyze multiple combinations of hemophore interactions, we used protein crosslinking as an alternative assay. Individual proteins and protein mixtures, 0.1 mM each, either without or with pre-incubation with an equimolar amount of hemin, were incubated with the crosslinker DSP and separated on acrylamide gels. Comparison without and with crosslinker indicated the appearance, after incubation with the crosslinker, of new species that correspond to the expected dimer size for several hemophores (Fig. S2). In order to test both homotypic and heterotypic interactions, we then incubated the Csa2 and Rbt5 (Fig. 4A) or Csa2 and Pga7 (Fig. 4B) hemophores, individually and together, in the apo or holo forms with the crosslinker, and separated the reaction products on an acrylamide gel (Pga7 and Rbt5 are too similar in size to be separated by SDS-PAGE). The results indicated that Csa2 homo-dimerization is unaffected by the presence of hemin, whereas Rbt5 and Pga7 dimerization is increased in the presence of hemin. Furthermore, Csa2-Rbt5 and Csa2-Pga7 heterodimers can be detected as well, and heterodimerization appears to also be increased in the presence of hemin.

**Fig. 4.**
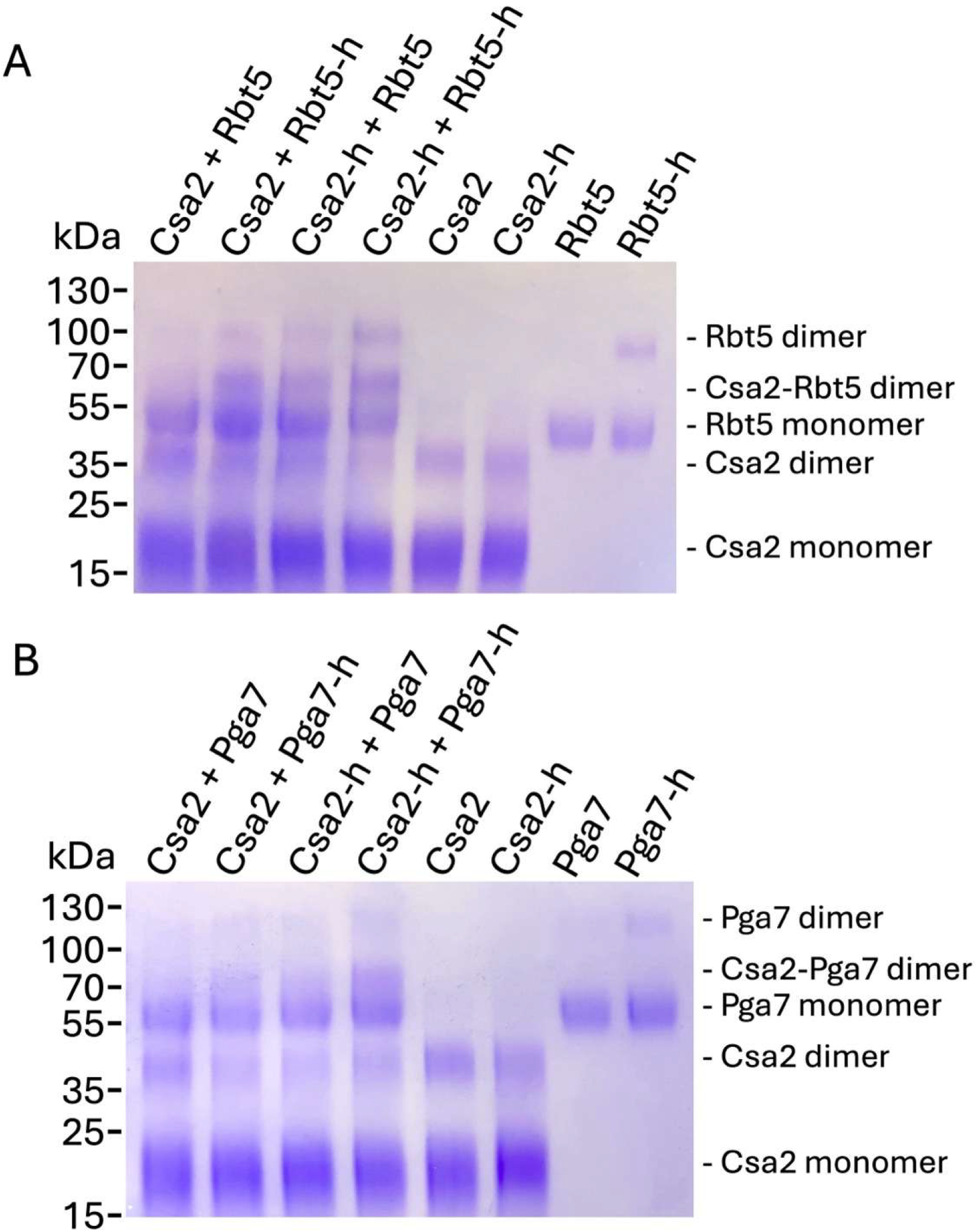
Crosslinking analysis of hemophore interactions. Incubation of Csa2 and Rbt5 (A) or Csa2 and Pga7 (B) with the crosslinker DSP cause the appearance of higher molecular-weight species corresponding to homodimers and heterodimers. Proteins (0.1 mM each) were incubated for 15 min at room temperature with the amine-reactive crosslinker DSP, and the reaction was then quenched with Tris-Cl. Reaction products were boiled in gel loading buffer without reducing agent, separated on a 4-20% gradient gel, and stained with Coomassie.

### Csa2 mutants defective in dimerization

To try to model the CFEM hemophore dimer interactions, we focused on Csa2, for which the monomer crystal structure had been determined (11). To predict the dimeric interaction site, we used five *in silico* docking softwares: Zdock, Patchdock, Swarmdock, ClusPro and Haddock (34–37). The vast majority of in silico docking predictions suggested that a specific face of the protein is involved in dimerization (Fig. S3). Two surface patches in particular, one including the residues Val86, Ile87, Pro88, and the second including Trp124 and Asp125, were found to participate in the dimer interaction in most predictions (Fig. 5A).

**Fig. 5.**
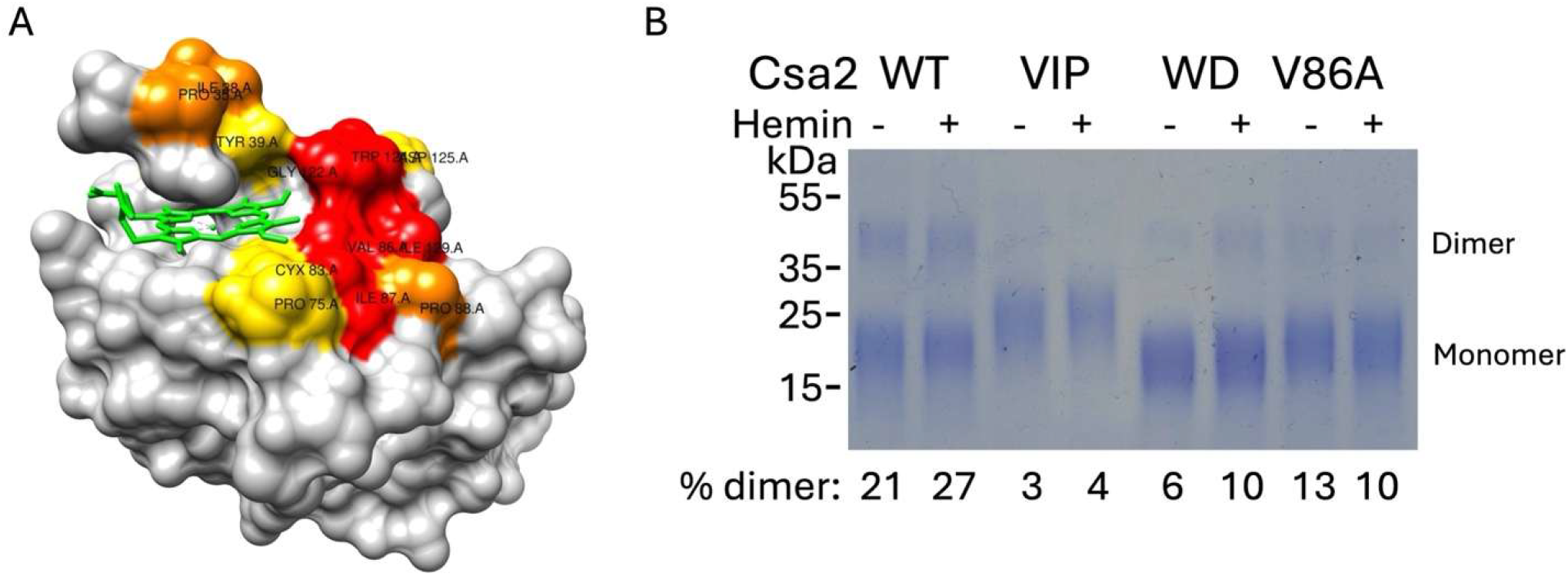
Csa2 dimerization mutants. (A) Residues predicted to most likely to participate in the dimer interaction in the were visualized by PyMol. (B) Csa2 wild-type and the indicated mutants were crosslinked and separated as in Fig. 4, and the Coomassie-stained gels were scanned and quantitated.

To test this quaternary structure model, the residues predicted to participate in dimerization were mutated to alanine, in clusters and individually, and their effect on dimerization was tested, using chemical crosslinking as a measure of interaction. As shown in Fig. 5B, whereas under our reaction conditions, approximately a quarter of the wild-type Csa2 protein was cross-linked in the dimer form, crosslinking went down to 12% for the V86A mutant, 7-11% for the W124A D125A mutant (WD), and to 3-4% for the V86A I87A P88A (VIP) mutants. The dimerization-defective mutants were nonetheless still able to bind heme based on isothermal calorimetry, with a calculated affinity similar to the wild-type affinity of 0.2 μM (11) and were also able to extract heme from hemoglobin immobilized on Sepharose beads (Fig. S4).

#### Heme exchange kinetics in Csa2 dimerization mutants

The Csa2 dimerization mutants V86A, WD and VIP were fused to GFP in order to monitor their heme exchange kinetics. We then measured the kinetics of heme transfer from the V86A and VIP mutant holo-proteins fused to GFP, to the wild-type native apo-Csa2. Transfer from these two mutants to the wild-type Csa2 showed somewhat reduced rates of transfer but maintained second-order kinetics. (Fig. 6). We next measured transfer between the mutant holo-proteins fused to GFP and their corresponding mutant apo-proteins. The single V86A mutant and the compound WD mutant showed substantially reduced reaction rates but maintained second-order kinetics. In contrast, the VIP mutant showed first-order kinetics, suggesting that in this mutant, protein-protein interaction was sufficiently reduced that it was not making a significant contribution to heme transfer kinetics.

**Fig. 6.**
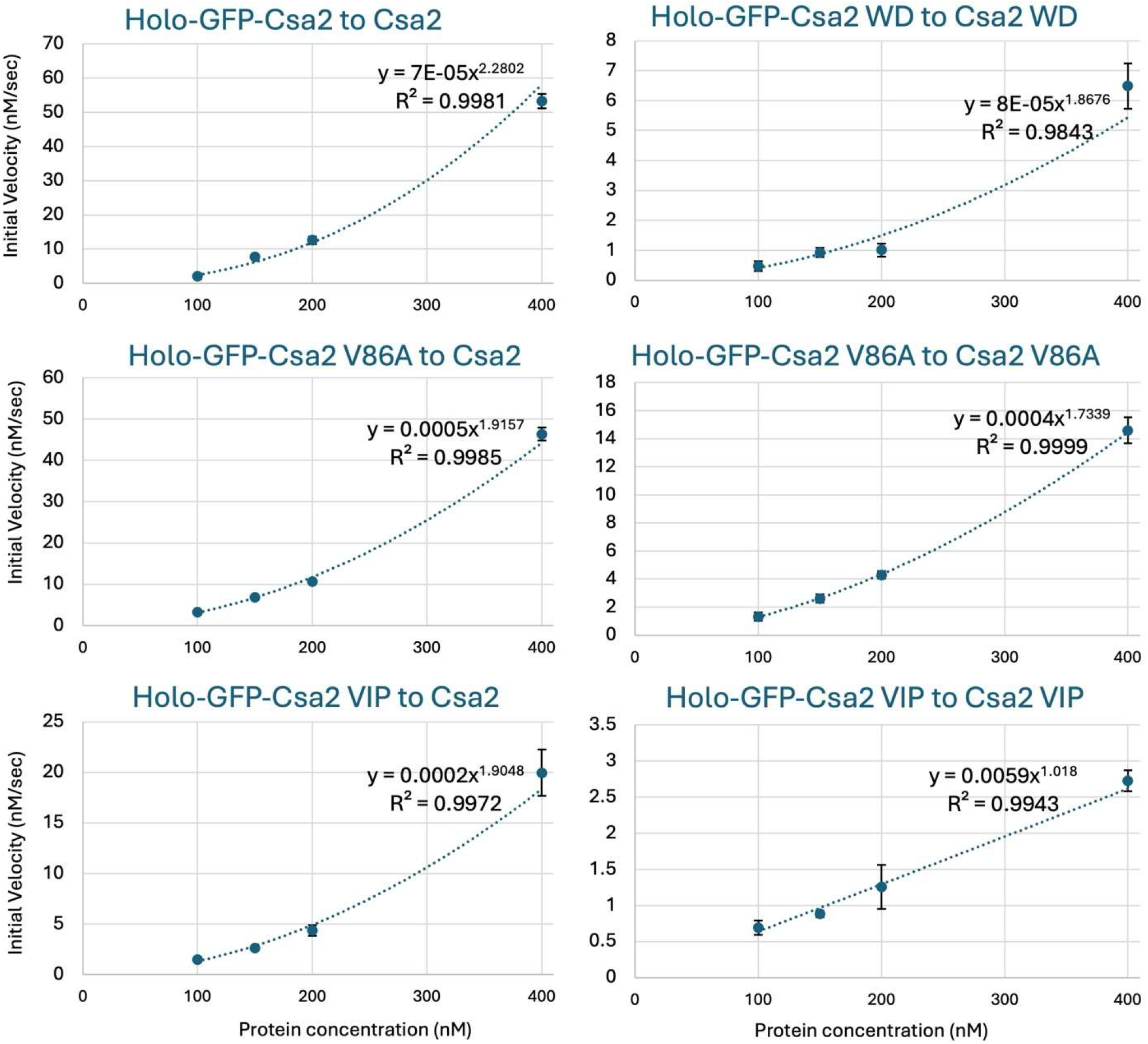
Heme exchange kinetics in the heme dimerization mutants. The indicated combinations of GFP-tagged wild-type and mutant holo-hemophores with the untagged native apo-hemophores were mixed, and de-quenching of the fluorescence by heme transfer was monitored with time at different starting protein concentrations, as in Fig. 3.

## DISCUSSION

Heme uptake is a common solution for iron acquisition in pathogenic microorganisms and, given the ubiquity of heme in nature, among saprophytic organisms as well (20). Like bacteria, fungi express different types of heme acquisition pathways, of which at least three were characterized, in *C. albicans, S. pombe* and *C. neoformans* (20, 38, 39). The *Candida spp*. heme uptake pathway includes a series of extracellular hemophores of the CFEM family that are able to extract heme from host proteins and to efficiently exchange it (11, 18, 19), resulting in a proposed pathway where the CFEM hemophores create a heme transfer cascade across the cell envelope (40). Missing from that model was the mechanism of transfer of heme to, and between, the CFEM hemophores.

In order to monitor the kinetics of heme binding to the hemophores, we relied on the observation that proximity of heme can quench the fluorescence of the green fluorescent protein EGFP (29–31). Early work indicated that linking a Cytochrome b_562_ domain to an EGFP domain resulted in a fusion protein that was quenched upon addition of hemin, but only up to about 70% (29), whereas later work indicated that up to 99% quenching could be obtained by embedding the cytochrome b_562_ domain within the EGFP domain (31). More recently, an attempt to analyze a *Corynebacterium diphtheriae* heme-binding domain by embedding it in EGFP also yielded a fusion protein that was quenchable by the addition of heme, but in this case the maximum quenching was 60%, possibly due to a greater distance between the heme binding site and the EGFP fluorophore than with the EGFP-Cytochrome b_562_ fusion (41). For the CFEM hemophores, we initially tried to fuse the EGFP domain adjacent to the hemophore domain, either N-terminally or C-terminally. The C-terminal fusions did reach up to 50% quenching with heme addition, but with the N-terminal fusions, over 90% quenching was obtained with Pga7 and Rbt5, and over 95% with Csa2. This indicates that embedding of a heme-binding domain within the EGFP domain is not always necessary for obtaining efficient fluorescence quenching, making FP-hemophore fusions potentially useful reagents for monitoring heme binding to the hemophores *in vivo*. We were also able to obtain quenching of the red fluorescent protein mScarlet fused to Pga7, which further expands the possibilities for use of these reagents *in vivo*.

As always when using FP fusions as reporter proteins, it is important to keep in mind the possibility that the added FP domain could alter the hemophore’s activity and interactions. We observed however that the fusion proteins can compete effectively with the native proteins for heme, suggesting that their activity is largely unaffected *in vitro*. Furthermore, we found in preliminary experiments (to be published separately) that at least some of the fusion proteins are active, fully or in part, when expressed in *C. albicans*, suggesting that they are similar in function to the native proteins.

The ability to monitor heme binding to the hemophores in real time enabled us to measure the kinetics of heme transfer from the host proteins to the fungal hemophores and between the hemophores, giving us potential insight into the reaction transfer mechanisms. Two basic heme transfer mechanisms can be considered: one where heme dissociates from one partner, and is captured by the second, without direct interaction between the proteins; and one where the two proteins interact before exchanging heme.

Reaction kinetics can in principle be used to derive the reaction mechanism: in the first mechanism, release-and-capture, where the reaction rate is determined by slow release of heme from a donor protein, the rate of transfer should increase linearly with concentration of the donor protein, i.e. show first-order kinetics. Alternatively, if the reaction depends on interaction between proteins, the rate of transfer can be expected to depend on the product of the concentrations of the two interacting molecules, i.e. would show second-order kinetics. However, in the classical enzyme kinetics calculations that predict the dependence of initial reaction velocities based on enzyme and substrate concentrations, the explicit assumption is that the enzyme-substrate complexes form a negligible part of the molecules pool, so that most of the interacting molecules are free in solution rather than in complexes (42). Where heme transfer between hemophores is concerned, if the fraction of hemophores that are in complexes becomes substantial, the amount of monomeric hemophores, which determine the reaction rate, will be reduced; in such a case, even though the reaction is bimolecular, the apparent order of reaction can be less than 2 (see Supplementary Methods). Another possibility for obtaining an apparent order of reaction between 1 and 2 is in the case of a mixed mechanism of release and capture together with some interaction-driven transfer; in this case however, the component of the reaction that depends on stochastic release of the heme will lower the overall reaction rate.

Heme transfer rates from hemoglobin to the hemophores were relatively slow and linear with concentration, consistent with first-order kinetics, suggesting that the mechanism of heme acquisition from hemoglobin involves stochastic release of heme from the globin, followed by capture of the heme by the CFEM hemophores. Interestingly, for heme bound to HSA, heme transfer rates were still low, but the reaction velocity dependence on concentration could be fitted to a power equation with an exponent slightly above 1 for the four types of FP-hemophores tested. This could indicate a mixed mechanism with partial dependence on interaction of the hemophores with the albumin, and might explain why the presence of HSA increases the ability of *C. albicans* to utilize hemoglobin heme (18).

In contrast to the host protein transfer kinetics, the heme exchange reactions between the three hemophores, in all combinations, was much faster and could be fitted to a power equation with exponents between 1.3 and 2, suggesting that the exchange reactions depend on protein-protein interactions. We tested this possibility by showing that the hemophores can be chemically crosslinked to form both homodimers and heterodimers. Interestingly, all interactions involving Rbt5 and Pga7 were increased in the presence of heme, but Csa2 homodimerization was not. However, the Csa2-Csa2 heme transfer rate was actually substantially faster than that of the other hemophores, and was pure second-order. It is possible that the stronger dimerization interactions of Pga7 and Rbt5 in the presence of heme, as detected by crosslinking, shift the reaction kinetics towards less-than second order due to a reduced monomer concentration; however, more quantitative measurement of the protein affinities will be needed to corroborate this explanation.

We probed the notion that the transient dimerization of Csa2 that we detected is responsible for the second-order reaction kinetics by constructing and testing Csa2 mutants defective in dimerization. Quaternary structure predictions of the Csa2 dimer pointed to a series of residues expected to participate in the dimeric interactions. Mutations of these residues resulted in mutants that dimerized less but were still able to bind heme and to extract it from hemoglobin. When reaction kinetics were measured, transfer between mutant and wild-type proteins maintained second-order kinetics and only weakly reduced rates. For the WD → AA and V86A mutants, transfer reaction where both the donor and acceptor are mutants also maintained second-order kinetics, however with much reduced rates of transfer, suggesting that dimerization still drove the reaction rate, but that the weaker dimerization resulted in slower exchange rates. Only the mutant most defective in dimerization, the triple mutant VIP → AAA, showed first-order kinetics of heme transfer in a homodimer and very low transfer rates, suggesting that the transfer in this mutant was governed by a stochastic release and recapture mechanism, similar to the mechanism of heme capture from hemoglobin.

Collectively, the second-order reaction kinetics (Fig. 3) and the DSP crosslinking experiments (Fig. 4), paint a picture of the importance of direct hemophore interactions in the heme transfer cascade of *C. albicans*, a notion confirmed by the mutagenesis data that link Csa2 dimerization to heme exchange kinetics (Figs. 5-6). We can now draw a more accurate model of the heme uptake mechanism of *Candida albicans* (Fig. 7) in which acquisition of heme from hemoglobin involves a release and capture mechanism of heme by the fungal hemophores and subsequently, a heme transfer cascade across the cell wall via hemophore-hemophore interactions.

**Fig. 7.**
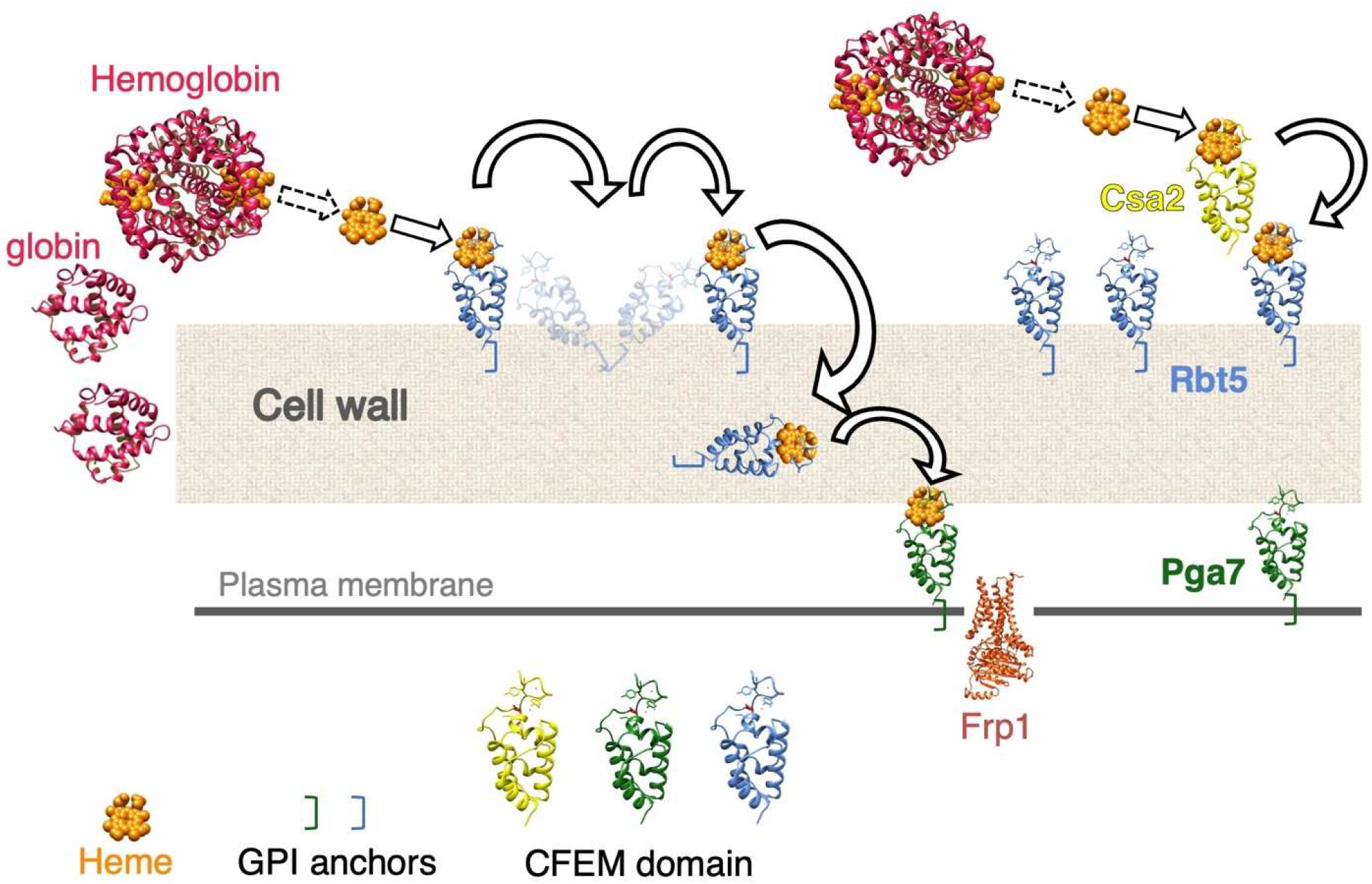
Model of the heme transfer reactions across the cell wall. See text for details.

One potential difficulty with this model is that Rbt5 and Pga7 are anchored to the cell wall matrix and the cell membrane, respectively, which could hamper protein-protein interactions. However, GPI-anchored cell wall proteins such as Rbt5 are attached to the end of extended, flexible β1,6 glucan chains (43). Combined with the abundance of Rbt5, estimated at 10^6^ molecules / cell (10) that suggests, based on a 100 μm^2^ surface area (44), an average horizontal distance of 10 nm between molecules, the β1,6 glucan chains chain attachment of these proteins may afford sufficient flexibility to enable protein-protein interactions within the cell wall matrix. The captured heme would eventually need to be delivered to the membrane-anchored Pga7 hemophore, which collaborates with the transmembrane protein Frp1 to effect its endocytosis (21).

To conclude, the development of heme-quenchable FP fusions with the fungal CFEM hemophores gave us the opportunity to address the mechanisms of heme acquisition and transfer. The analysis of heme exchange kinetics with the fluorescent hemophores enabled us to identify and characterize protein-protein interactions between CFEM hemophores as being important for heme exchange, and to refine our understanding of the mechanism of host heme acquisition by fungal pathogens. FP-hemophore fusions represent a new tool in the study of heme transport that carry the potential to illuminate other aspects of this pathway in *C. albicans* and other microorganisms.

## METHODS

### Strains and plasmids

All protein expression was done in *Pichia pastoris* X-33 (Invitrogen) transformed with the relevant plasmids according to the Invitrogen manual.

### Plasmid construction

*P. pastoris* expression plasmids were based on pICZαA plasmids expressing Rbt5, Pga7 and Csa2 (11, 19) with the addition of *Candida*-optimized GFP and mScarlet (mSca) genes (32, 33) by Gibson assembly, at the positions indicated in Table 1. Mutations were introduced by site-directed mutagenesis.

**Table 1.** List of plasmids for *P. pastoris* expression.

|  |  |  |
| --- | --- | --- |
| pICZαA | ZEO<sup>R</sup>, AOXp, 6xHis, Myc epitope, a-factor secretion signal | Invitrogen |
| KB2366 | Csa2 (a.a.19-147)-Myc-6xHis | Nasser et al.<br>(2016) |
| KB2520 | Csa2 (a.a.19-147)V86A I87A P88A-Myc-6xHis | This work |
| KB2521 | Csa2 (a.a.19-147)W124A D125A-Myc-6xHis | This work |
| KB2535 | Csa2 (a.a.19-147)V86A-Myc-6xHis | This work |
| KB2807 | GFP-PGA7 (a.a. 17-195)-Myc-6xHis | This work |
| KB2808 | PGA7(a.a. 17-195) $\nabla$145-GFP | This work |
| KB2809 | mScarlet GFP-PGA7 (a.a. 17-195)-Myc-6xHis | This work |
| KB2810 | GFP-RBT5 (a.a. 22-219)-Myc-6xHis | This work |
| KB2811 | RBT5 (a.a. 22-219) $\nabla$147-GFP | This work |
| KB2834 | GFP-CSA2(a.a. 33-147) | This work |
| KB2841 | GFP-CSA2(a.a. 33-147) (VIP) | This work |
| KB2842 | GFP-CSA2(a.a. 33-147) (WD) | This work |
| KB2843 | GFP-CSA2(a.a. 33-147) (V86A) | This work |

### Hemin and hemoglobin preparation

All stocks were freshly prepared before each assay. Stock solutions were 0.5 mM bovine hemoglobin (methemoglobin, i.e. ferric hemoglobin) or 0.5 mM human serum albumin (Sigma) in PBS, or 2 mM hemin (Frontier Scientific) in 10 mM NaOH.

### Recombinant protein production

CFEM hemophores and FP-hemophore fusions: the various wild-type and mutant native and FP-fused hemophores were expressed as C-terminal 6xHis-tagged proteins in the *P. pastoris* expression system as described (Pichia expression manual, Invitrogen, Carlsbad, CA). Briefly, cells from an overnight culture in BMGY were diluted in BMMY to OD_600_=1. Cultures were then incubated at 28°C with vigorous shaking for 120 hours, with addition of 0.5% methanol every 24 hours. The cells were removed by centrifugation (4500 rpm, 5 min), and the pH was adjusted to 8.0 with 5 M KOH. The medium was centrifuged again (5400 rpm, 5 min in 50 ml Falcon tubes), and then the supernatant was transferred to a glass cylinder and incubated with nickel beads with shaking at RT for two hours. After two hours the cylinders were left to stand on the bench overnight to let the beads sediment. The supernatant was carefully transferred with a pipette to 50 ml Falcon tubes for rinsing the beads. Four washes were made in pH=8 with different concentrations of imidazole, 10 mM for first wash, 20 mM for the second and third wash and 50 mM for the fourth wash. Then protein was then eluted with 250 mM imidazole. The proteins were then concentrated to 500 μL with a Centricon filter, and to remove bound heme from the medium, 3 M imidazole was added, and the tubes were tumbled overnight at 4°C. The heme precipitate was spun down for 10 min in an Eppendorf centrifuge and the apo-protein supernatant was transferred to a dialysis tube and dialyzed against a 1000-fold volume of PBS once for 2 h at room temperature, and a second time overnight at 4°C. Concentration was then determined by absorbance, based on a predicted absorbance coefficient calculated from the protein sequence.

Holo-FP-hemophores: to generate heme-saturated hemophore stocks, the apo-FP-hemophores were mixed with a 50% excess amount of hemin chloride. Excess free hemin was then removed either by passing the protein over a size exclusion chromatography column (GE Superdex 75 10/300) or by diluting the mix and concentrating > 10x three times with a Centricon filter.

### Reaction kinetics measurements

For equilibrium fluorescence measurements, a Tecan Infinite M Plex plate reader was used. For kinetics experiments, fluorescence was monitored with an Agilent Cary Eclipse spectrofluorometer equipped with a TgK Scientific SFA-20M microvolume stop-flow injector driven by an OPT-20P pneumatic drive and a quartz cuvette with a 2 / 10 mm path lengths (excitation / emission). Measurements were taken typically every 0.05 - 0.2 sec, depending on reaction speed. GFP was measured at 440 nm excitation, 508 nm emission wavelengths, and mScarlet at 550 nm and 596 nm, respectively, all with a 5 nm slit and PMT sensitivity set at 600 V. Velocity was measured in the initial, more linear stage of the reaction, corresponding to the first 10% change compared to the fluorescence change at equilibrium. Hemophore protein concentration was measured by absorbance with a Cary 60 spectrophotometer, using the calculated specific absorbance based on protein sequence. Five to eight injections were measured per concentration point. Fluorescence units were converted to concentration based on initial fluorescence for the apo-FP-hemophores, or on fluorescence after heme removal by treatment with 2 M imidazole for the holo-FP-hemophore preparations.

### Protein crosslinking

Purified proteins were diluted to 0.1 mM each, with or without 0.1 mM hemin, together with 15 mM dithiobis[succinimidylpropionate] (DSP; Thermo-Scientific) in crosslinking buffer (20 mM Na-HEPES pH 7.5, 100 mM NaCl, 1 mM EDTA, 1.5 mM MgCl_2_) and incubated for 15 min at room temperature. The reaction was quenched by addition of 20 mM Tris pH 7.5 and incubated a further 15 min. The samples were then boiled with protein loading buffer without reducing agent, separated on a 4%-20% SDS-PAGE gradient gel (BioRad) and stained with Coomassie. Scanned gels were quantitated with the TotalLab software.

### Isothermal titration calorimetry (ITC)

Titrations were performed at 25 °C using a MicroCal iTC200 system (GE Healthcare). The protein stocks and a freshly prepared hemin stock solution were diluted in PBS. All the injections were carried out at 150 second intervals. To prevent/minimize heme adsorption, the calorimeter cell and the microsyringe used for the injections were washed extensively with 10 N NaOH after each experiment. Protein concentration was determined using a NanoDrop spectrophotometer, and hemin stock concentration was determined by absorbance at 398 nm in an organic solution ^(45)^. For the titration experiments, the concentrations of protein were 140 μM in the syringe and 20 μM hemin in the cell. The resulting titration data were analysed and fitted using the Origin for ITC software package supplied by MicroCal to obtain the stoichiometry (n), the dissociation constants (K_D_) and the enthalpy (ΔH) and entropy (ΔS) changes of binding. For the fit, any constraints on the stoichiometry and ΔH were not fixed.

## ACKNOWLEDGMENTS

We thank Fabian Glazer (Technion Bioinformatics Knowledge Unit) for help with visualization of the Csa2 dimer interacting face, Rob Arkowitz (U. Nice) and Jamie Konopka (SUNY Stony Brook) for *Candida*-adapted FP plasmids, and Abdussalam Azem (Tel Aviv university), Amit Reddi (Georgia Tech) and Amnon Horovitz (Weitzmann Institute) for discussions. This work was supported by grants 587/19 and 1969/24 from the Israel Science Foundation.

## SUPPLEMENTARY FIGURES

**Fig. S1.**
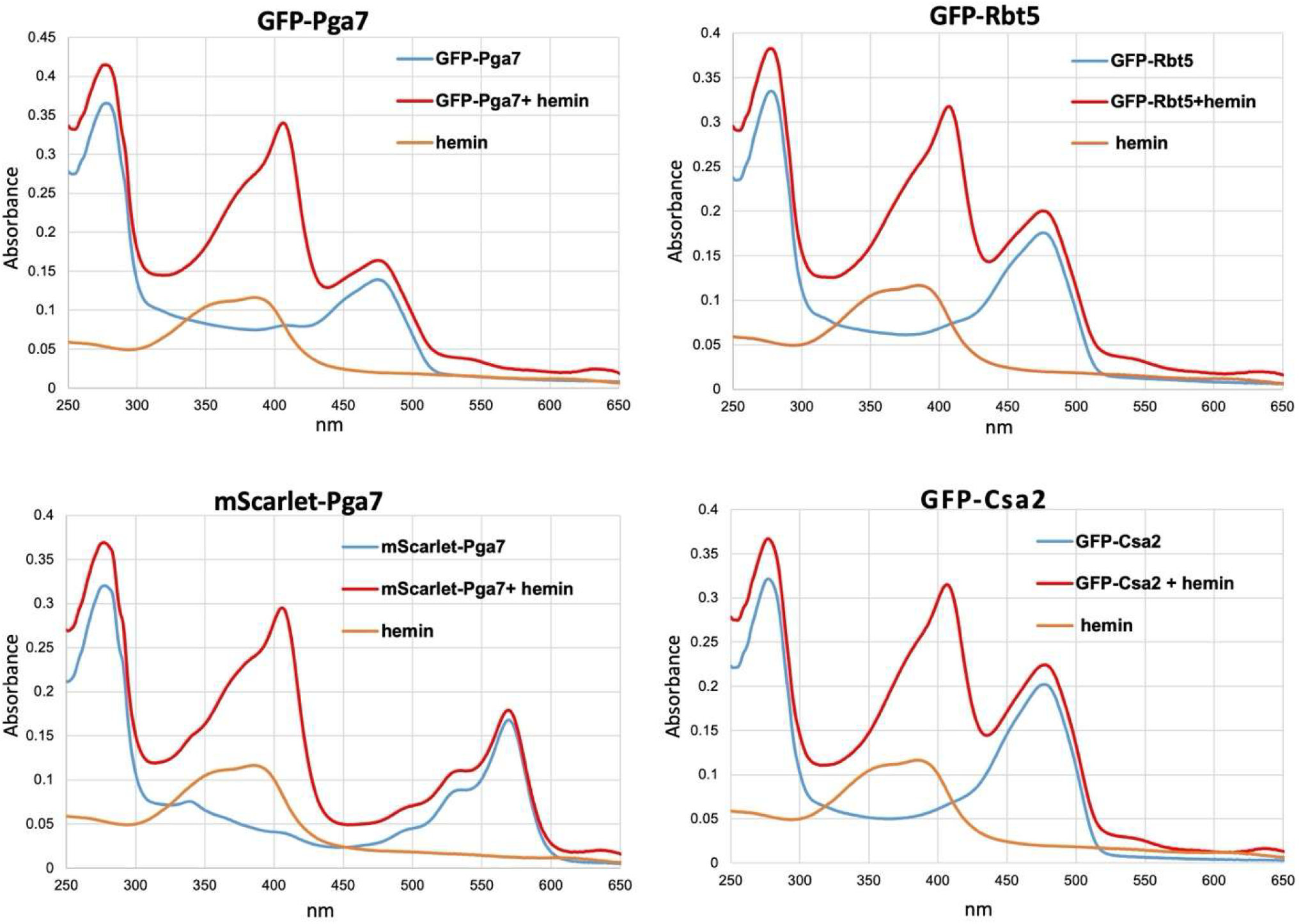
Absorbance scans of fluorescent protein - CFEM hemophore fusions with and without heme. Absorbance spectra of the indicated proteins (5 μM each), alone or mixed with 3 μM hemin. The 406 nm absorbance peak is typical of heme binding to the CFEM hemophores.

**Fig. S2.**
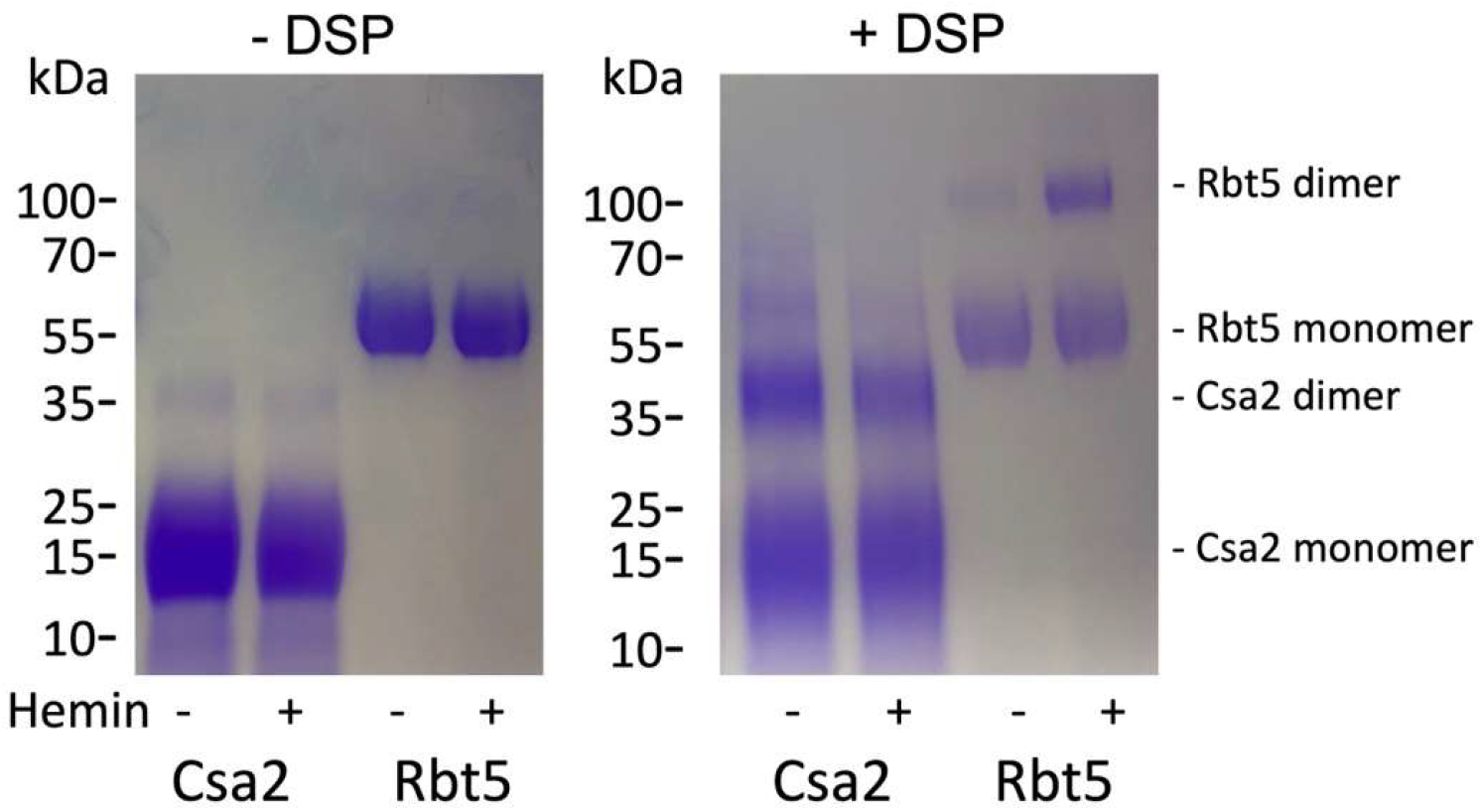
Crosslinking analysis of hemophore interactions. Incubation of Csa2 and Rbt5 with the crosslinker DSP cause the appearance of higher molecular-weight species corresponding to a dimer weight.

**Fig. S3.**
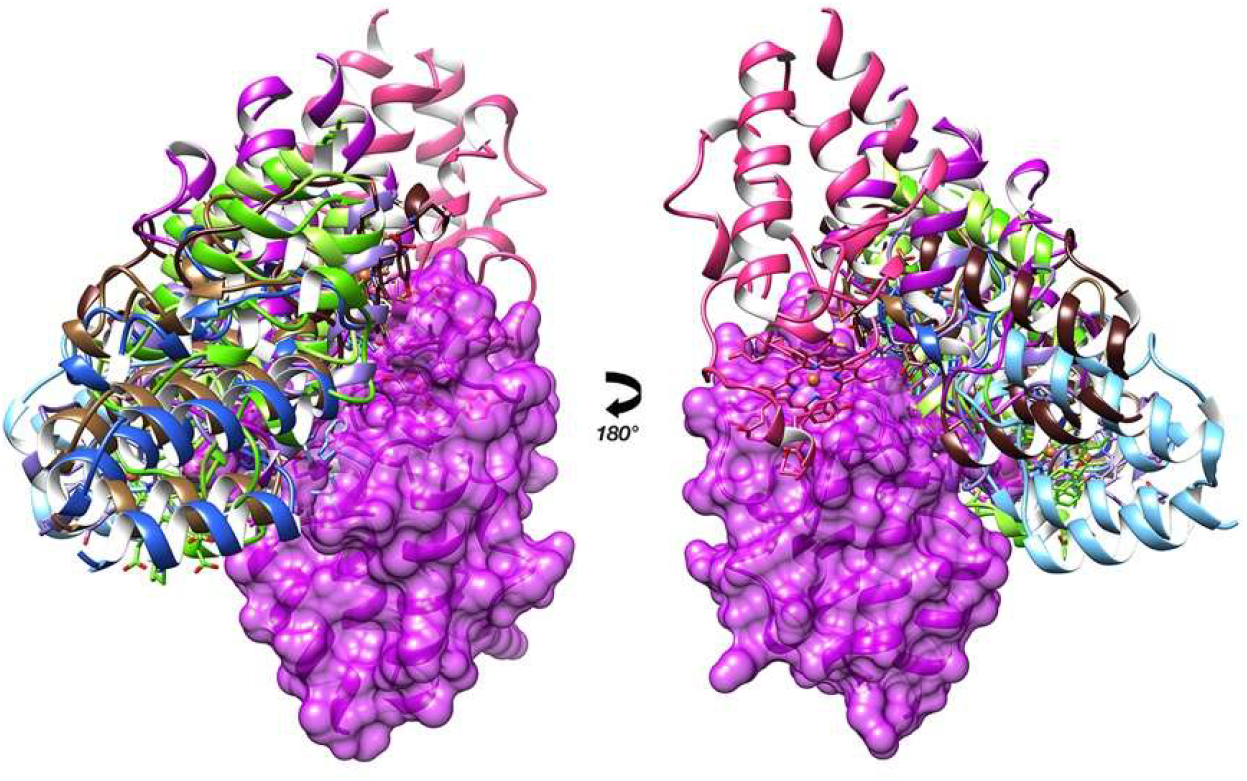
Csa2-Csa2 dimer structure predictions. The ten best Firedock docking predictions are shown as Csa2 ribbon models docked against a fixed solid surface Csa2 monomer.

**Fig. S4.**
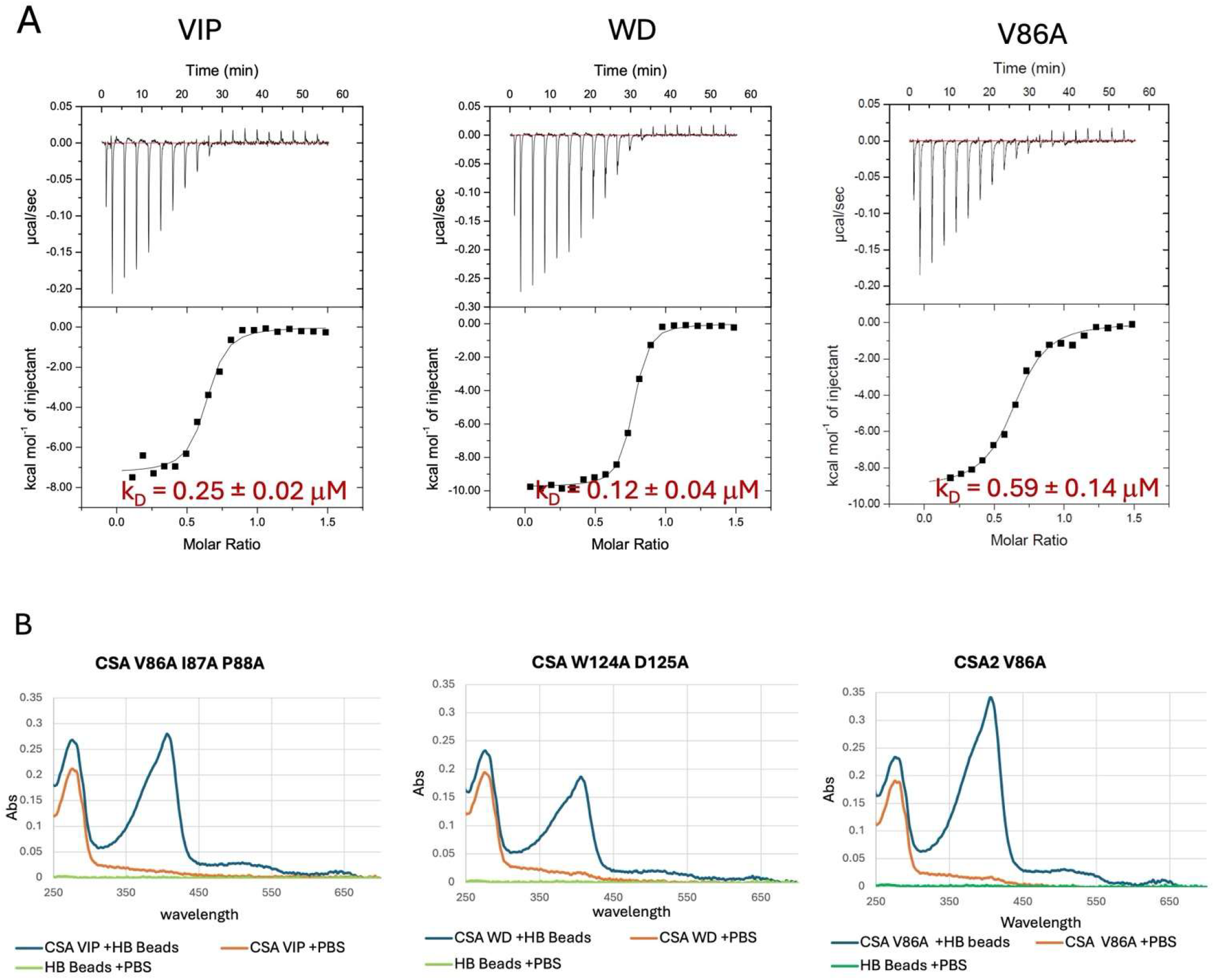
Heme binding in the Csa2 dimerization mutants. A. Isothermal calorimetry of heme binding to the indicated dimerization mutants. 150 μM protein was gradually injected into a chamber with 20 μM hemin. The calculated affinity in red is the average of three experiments. B. Heme extraction from hemoglobin beads by the Csa2 dimerization mutants. Hemoglobin covalently conjugated to CnBr sepharose beads was incubated with the indicated Csa2 mutants, and the Csa2-heme complex was detected in the supernatant as a 406 nm Soret absorption peak by UV/VIS absorbance spectroscopy.

## Supplementary Methods

Prediction of the dependence of transfer rates of heme from hemophore H1 to H2 on the concentrations of holo-H1 and apo-H2: we want to calculate the rate of production of the free GFP-CFEM hemophore H_1_, where H_1_h is the initially heme-loaded GFP-CFEM hemophore and H_2_ is the initially heme-free native hemophore. We assume that fluorescence de-quenching occurs upon release of H_1_ from the complex.

Let

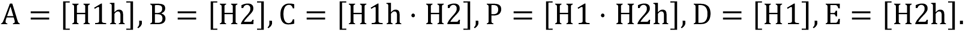

Assuming that the reaction is fully symmetrical, a minimal mechanism is

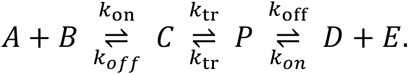

*k*_*on*_, *k*_*off*_: formation and breakup of the protein complexes H1h⋅H2 and H1⋅H2h

*k*_*tr*_: intra-complex ligand transfer is symmetrical based on equal distribution of heme between CFEM hemophores (11, 19).

We will calculate for the case where the two starting species are equimolar, A = B = R_0_

Assume the reaction rapidly reaches a quasi-steady-state so that

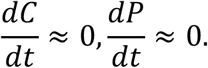

Using a quasi-steady-state treatment for *C*and *P*, respectively, early in the reaction, when [D] and [E] are negligible:

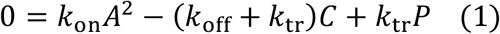

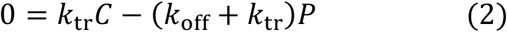

Rearranging (2) gives

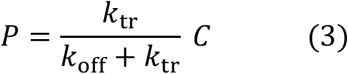

By substituting P from equation 3 into equation 1, one obtains

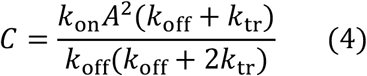

Substituting C in equation 3 using 4, one obtains

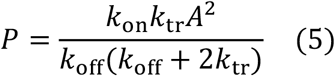

Since at early time points, when [D] and [E] are low, the starting velocity *v*_0_ will be given by

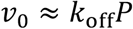

Then, substituting P from (5):

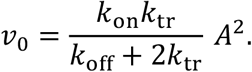

So the rate is always proportional to the **free** reactant concentration squared.

### Relation of free concentration *A* to *R*_0_

Because each complex consumes one *A* and one *B*, early on

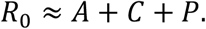

From combining the steady-state relations (1) and (2) and rearranging, one obtains:

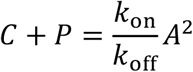

If

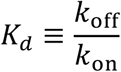

Hence

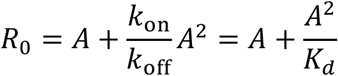

That equation determines the crossover between first and second order.

### When is the transfer reaction expected to be second order?

If binding is weak or concentrations are low, so that most material remains free:

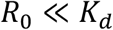

then *A* ≈ *R*_0_, and

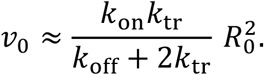

So the observed initial rate is **second order** in starting reactant concentration.

### When is the transfer reaction expected to be first order?

If binding is strong or concentrations are high, so that most material is tied up in *C* + *P*:

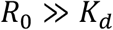

then from

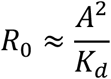

we get

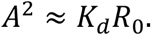

Substitute into the rate law:

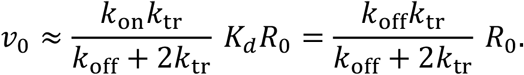

So the observed initial rate is **first order** in starting reactant concentration.

### Summary

Let

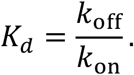

Then:

- **Second-order regime** when

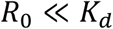

because *A*_free_ ≈ *R*_0_, so 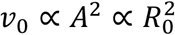.
- **First-order regime** when

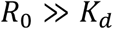

because most reactant is already in *C* + *P*, so 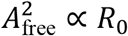, giving *v*_0_ ∝ *R*_0_.

When the starting concentration is close to the hemophore complex dissociation constant *K*_*d*_, the apparent reaction order will be between first and second order.

